# From Pairwise Distances to Neighborhood Preservation: Benchmarking Dimensionality Reduction Algorithms for CyTOF, scRNA-seq, and CITE-seq

**DOI:** 10.1101/2025.04.28.651069

**Authors:** Polina Bombina, Zachary B. Abrams, Reginald L. McGee, Kevin R. Coombes

## Abstract

Dimensionality reduction algorithms are essential tools for visualizing high-dimensional biological data, such as single-cell transcriptomics, mass cytometry by time of flight, and cellular indexing of transcriptomes and epitopes. These algorithms map complex, high-dimensional data into lower dimensions to reveal underlying structures and patterns. Despite the popularity of dimensionality reduction methods like t-SNE and UMAP, concerns have arisen regarding their ability to preserve critical aspects of high-dimensional data and their sensitivity to user-defined parameters. This study aims to evaluate the impact of extreme dimension reduction from hundreds or thousands of dimensions to just two dimensions, highlighting the resulting distortions to deeply understand their implications. Given the significance of dimensionality reduction in biological research, careful evaluation of these methods is necessary to ensure reliable and meaningful results. In this paper, we present a comprehensive evaluation of 16 dimensionality reduction methods. Our evaluation addresses several key factors, such as the preservation of pairwise distances and local neighborhood relationships between the original high-dimensional space and the low-dimensional projections.

## 1 Introduction

Dimensionality reduction (DR) algorithms have become essential tools for researchers to explore high-dimensional biological data. These algorithms transform complex, multidimensional data into a lower-dimensional representation, facilitating visualization and analysis. Due to the complexity and volume of data, DR methods are indispensable for other analyses, such as clustering, classification, pathway discovery, and cell type identification.

Different DR methods serve distinct purposes. Some methods, like Principal Component Analysis (PCA) [1], aim to preserve global structures by maximizing the variance explained, while others, such as t-Stochastic Neighbor Embedding (TSNE) and Uniform Manifold Approximation and Projection (UMAP),[2,3] focus on maintaining local relationships between data points. Techniques like Isomap [4] and Multidimensional Scaling (MDS) [5] emphasize the preservation of geodesic and pairwise distances, respectively. Additionally, methods like Independent Component Analysis (ICA) [6] and Locally Linear Embedding (LLE) [7] seek to capture more complex, non-linear structures within the data. Each approach is tailored to reveal different aspects of the data’s underlying patterns, making the choice of method crucial for achieving meaningful insights.

Several earlier studies have benchmarked DR algorithms to assess their stability, accuracy, and computational efficiency across different biological datasets. Xiang et al. [8] compared ten widely used DR techniques—PCA, ICA, Zero Inflated Factor Analysis (ZIFA) [9], GrandPrix [10], TSNE, UMAP, scvis [11], VAE [12], and SIMLR [13]—evaluating their performance in preserving data structure using clustering-based metrics such as Adjusted Rand Index (ARI) [14], Normalized Mutual Information [15], and Silhouette Width [16]. These methods were applied to both simulated and real single-cell transcriptomics (scRNA) datasets, providing insights into their utility for biological data analysis.

Building on previous work, Huang et al. [17] introduced two novel approaches—Local Supervised Evaluation and Local Unsupervised Evaluation—that refined the assessment of DR techniques. Local Supervised Evaluation tests how well DR methods preserve local structure using labeled datasets and supervised algorithms, such as support vector machines (SVM) and k-nearest neighbors. In contrast, Local Unsupervised Evaluation is applied to unlabeled data and focuses on preserving neighborhood relationships between high- and low-dimensional representations. This approach uses metrics like Random Triplet Accuracy [18], Distance Spearman Correlation [19], and Centroid Distance Correlation (introduced by Huang et al.) to provide further insights into data preservation during DR on scRNA data.

Wang et al. [20] benchmarked 21 DR techniques on both real and synthetic mass cytometry (CyTOF) samples. They introduced the Point-Cluster Distance metric, which computes Euclidean distances between cells and cluster centroids, to handle memory-intensive pairwise distance calculations. Their study included multiple evaluation metrics such as Spearman’s Correlation, Earth Mover’s Distance (EMD) [21], Neighborhood Proportion Error [22], and ARI to assess the preservation of local structure and clustering performance across different DR methods.

Cooley et al. [23] introduced the average Jaccard distance metric for evaluating DR techniques based on the preservation of local neighborhoods. Their approach compares the local neighborhood of a point in the original high-dimensional space with its representation in the reduced-dimensional space using Jaccard distance. This method, applied to techniques such as TSNE, Isomap, UMAP, and MDS, provides new perspectives on distortions introduced during DR, offering a more nuanced understanding of how these methods perform on synthetic and real biological datasets, including scRNA-seq.

Despite their widespread use, methods like TSNE and UMAP are not without limitations. These techniques often depend heavily on user-defined parameters, such as perplexity in t-SNE and minimum distance in UMAP, which can significantly alter the resulting embedding. Moreover, while these methods excel at preserving local structure, they can distort global relationships between clusters, potentially leading to misleading biological interpretations. Given their central role in biological analyses, it is crucial to systematically evaluate how such methods perform in preserving both local and global data structures under extreme dimension reduction.

The present study uniquely benchmarks DR methods across several major types of biological data: including scRNA, CyTOF, and cellular indexing of transcriptomes and epitopes (CITE-seq), filling a critical gap in existing research. No previous study has comprehensively evaluated DR techniques across a diverse array of biological datasets using a unified approach. Our study will employ advanced distortion metrics to assess the performance of various DR methods in preserving both global and local data structures. Notably, we will refrain from using clustering quality metrics in this evaluation. Without access to ground truth, clustering quality metrics may not accurately reflect the true effectiveness of DR, making them less relevant for our analysis.

## 2 Materials and Methods

### 2.1 Terminology

Both developers and evaluators of DR algorithms have used a variety of terms for the lower-dimensional results of applying these algorithms: projections, mappings, embeddings, reduced spaces, and others. In this paper, we will refer to all of these results as “projections”.

### 2.2 Dimensionality Reduction Algorithms

We used the Rdimtools R package [24], version 1.1.2, which supports a comprehensive suite of 145 DR algorithms, including 109 unsupervised methods. To narrow this extensive collection down to the most appropriate methods for our study, we followed a systematic approach involving data simulation and visual evaluation.

We simulated data points on two geometric shapes: a sphere and a torus. For both the sphere and the torus, we simulated 900 random 3D points uniformly distributed on the surface of each shape, and we used six arbitrary clusters produced by hierarchical clustering with the `Mercator` R package (version 1.1.4) to aid visualization.

We applied each algorithm to these datasets using default parameters to generate two-dimensional projections. The resulting projections were visualized, and plots were assessed for their ability to preserve the expected structure of the original shapes. Algorithms producing projections that appeared reasonable and interpretable were chosen for further analysis. when multiple algorithms gave very similar results on both simulated data sets we only retained a small number of representtive algorithms, to avoid redundancy. Additionally, we included TSNE and UMAP because they are widely recognized and commonly used methods in biology and bioinformatics. The UMAP analysis was conducted using the `umap` R package (version 0.2.10.0) [25], since UMAP is not included in `Rdimtools`

Using this procedure, we selected 16 algorithms for this study (Table 1). Fig. 1 and Fig. 2 below illustrate the projections of spherical and torus data for these 16 algorithms.

**Table 1:**
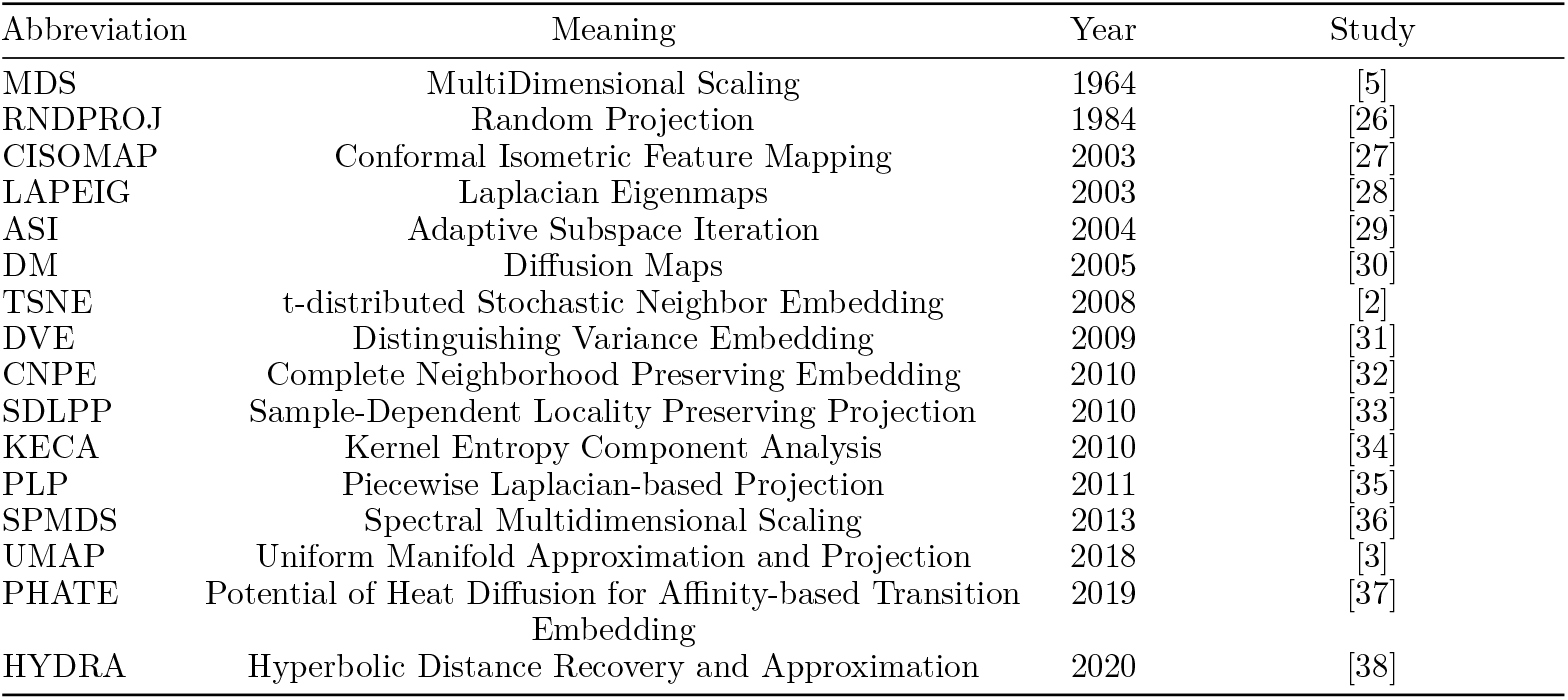
List of selected dimensionality reduction algorithms (chronological order).

**Figure 1.**
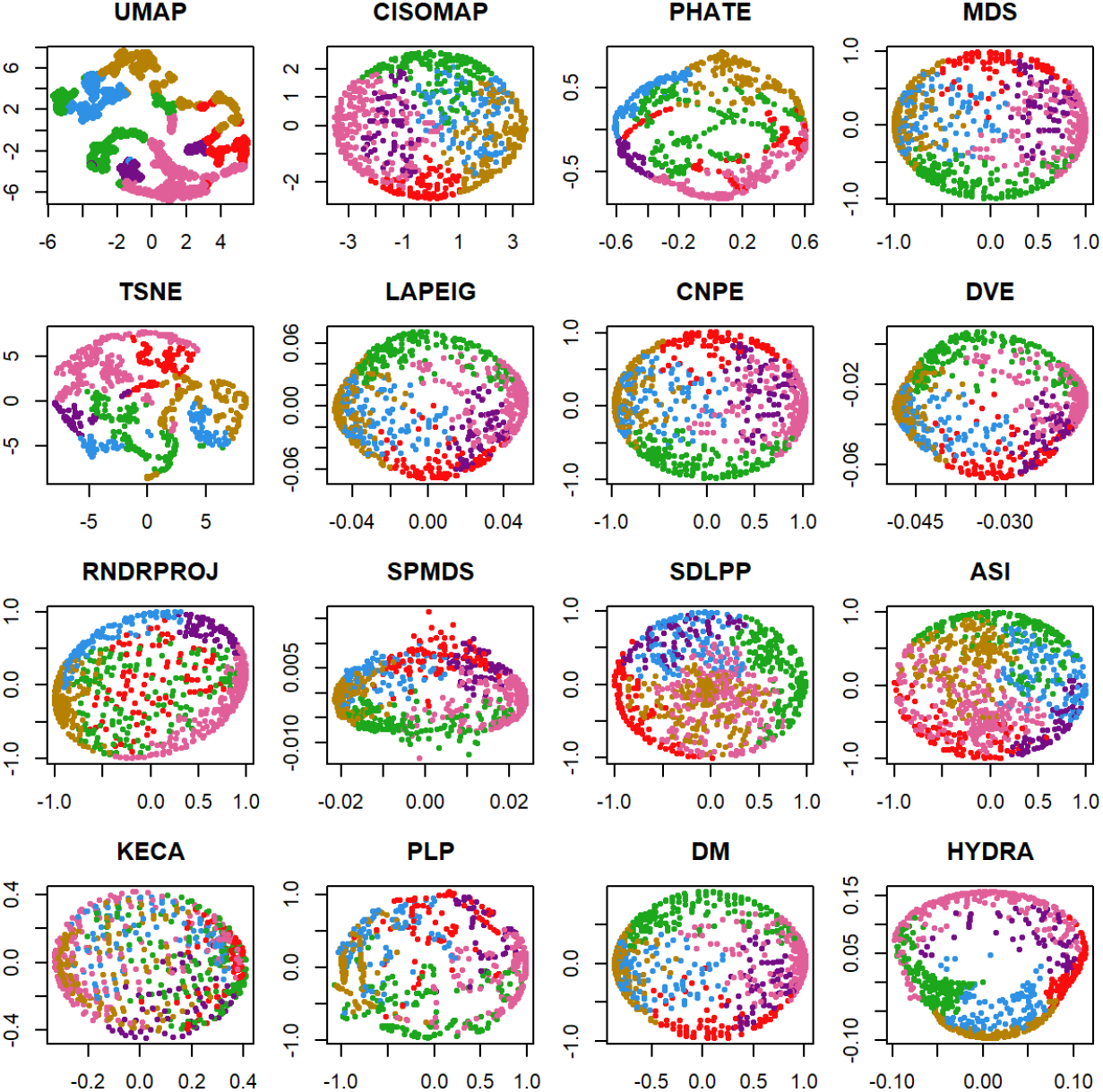
Projections of spherical data using sixteen DR algorithms.

**Figure 2.**
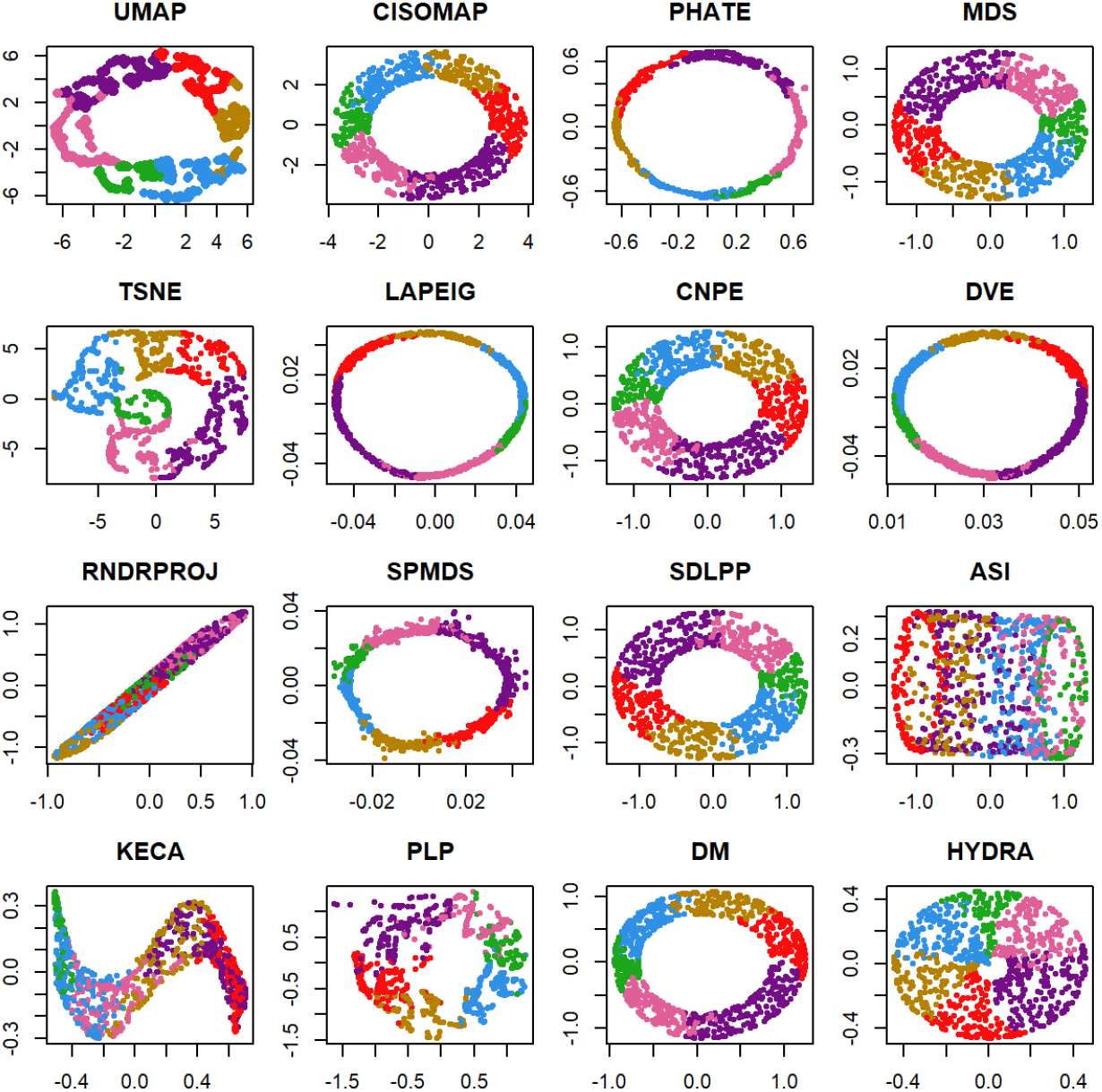
Projections of a torus using sixteen DR algorithms.

While we do not delve into the mathematical background of these methods here, readers interested in the technical details are encouraged to consult the relevant documentation and references for further insights.

### 2.3 Quality Metrics for Dimensionality Reduction Algorithms

From the wide range of evaluation metrics reviewed above, we focused on quality metrics that do not rely on known labels or clusters (ARI, NMI, and Silhouette Width were omitted). We classify the quality metrics into two main categories depending on whether they are based on *Pairwise Distance Matrices* or *Neighborhood Preservation*. These metrics evaluate different aspects of the preservation of pairwise distances, local neighborhood relationships, and topological structure between the original high-dimensional space and the low-dimensional projections.

We use the following notation to briefly describe the metrics. Let *A* ∈ ℝ^*n*×*M*^ be a high-dimensional data matrix, where rows are data points 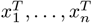 and *Ã* ∈ ℝ^*n×m*^ be a low-dimensional reduction matrix. *d*_*ij*_ *d*_*ij*_ denote pairwise distances between points *i* and *j* 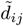 and in high- and low-dimensional spaces, respectively.

#### 2.3.1 Metrics Based on Pairwise Distance Matrices

##### Milnor’s Distortion

Milnor’s method [39] was initially used to measure changes to the “scale” of a mapping between points on a spherical earth and their projections onto a map in the Euclidean plane. However, the definitions generalize naturally to arbitrary DR algorithms to quantify the distortion in the projection, defined as the ratio of the maximum to the minimum scale. Let *d*_*S*_(*x*_1_, *x*_2_) denote the distance between two points *x*_1_ and *x*_2_ in a high-dimensional space, and let *d*_*E*_(*f* (*x*_1_), *f* (*x*_2_)) be the distance between their images under a projection *f* to a lower-dimensional Euclidean space. The scale of the map with respect to points *x*_1_ and *x*_2_ is defined as the ratio *d*_*E*_(*f* (*x*_1_), *f* (*x*_2_))*/d*_*S*_(*x*_1_, *x*_2_). Now let Let *σ*_1_ be the minimum scale as *x* and *y* vary and let *σ*_2_ be the maximum scale. The Milnor distortion of the projection *f* is defined to be

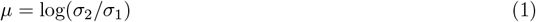

##### Sigma Distortion

The *σ*-distortion metric [40] quantifies how well distances between data points are preserved in the low-dimensional space. It evaluates the normalized ratio of distances for each point pair, comparing how well the distances in the low-dimensional representation match the original high-dimensional distances. Let *X*_n_ be a finite subset of *X*. Given a distribution *P* over *X*, let ∏ =*P* × *P* denote the distribution on the product space *X X*. For any projection *f*, let 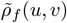 denote the normalized ratio of distances given by

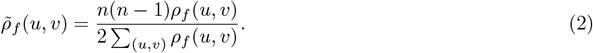

Here, *ρ*_*f*_ (*u, v*) = *d*_*Y*_ (*f* (*u*), *f* (*v*))*/d*_*X*_ (*u, v*) (if close to 1, then distortion is small). The *σ*-distortion is defined as:

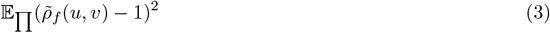

##### Stress

Stress is a well-known metric used to assess the quality of a projection [41]. It calculates the root mean square of the difference between the pairwise distances in the high-dimensional and low-dimensional spaces. Minimizing stress guarantees that the relative distances between points are preserved as accurately as possible. Stress is defined by:

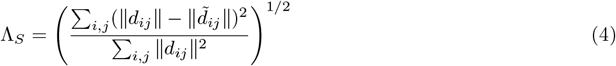

##### Spearman’s Rho

This metric is defined as the correlation between the vector distances in the original high-dimensional space and that in the low-dimensional space.

##### M1 (Mean-Squared Distortion)

*M* 1 [41] measures the distortion of pairwise distances by calculating the mean squared difference between the distances in the high-dimensional space and those in the low-dimensional space. This metric is useful for assessing the retention of “total variance” during dimensionality reduction.

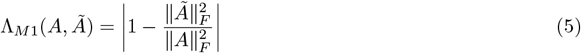

##### Earth Mover’s Distance

Earth Mover’s Distance measures the structural difference between two distributions, treating the problem as the work required to transform one distribution into another. It provides a way to assess the geometric similarity between the original and reduced representations of data points.

##### Implementation Notes

We developed our own custom functions to compute Milnor’s distortion, Sigma distortion, Stress, and M1 metrics. To compute Spearman’s Rho, we used the `SpearmansRho` function from version 0.2.1 of the `DRquality` R package [42]. We used the `emd` function from the `emdist` R package to calculate Earth Mover’s Distance.

#### 2.3.2 Metrics Based on Neighborhood Preservation

##### Average Jaccard Distance

This metric, based on the intersection and union of sets of nearest neighbors, calculates the similarity between the neighborhoods of points in the original high-dimensional space and the low-dimensional projection [43]. For each data point, the neighborhood consisting of the nearest *k*-neighbors were found in the ambient space, call this set *A*, and the projected space, call this set *B*. To calculate the Jaccard distance between *A* and *B*, use the usual definition

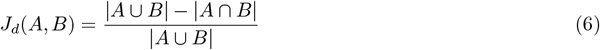

The Average Jaccard Distance was calculated by taking the arithmetic mean of the Jaccard distance over all points.

##### Trustworthiness

Trustworthiness [44] identifies how well the low-dimensional representation preserves the nearest neighbors of each data point from the high-dimensional space. A higher trustworthiness indicates better preservation of local structure. Calculate trustworthiness using the following definition:

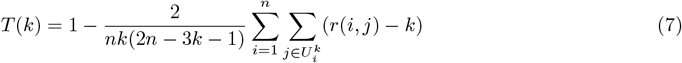

Here, *r*(*i, j*) represents the rank of the low-dimensional datapoint *j* according to the pairwise distances between the low-dimensional data points. 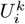 indicates the set of points that are among the *k* nearest neighbors in the low-dimensional space but not in the high-dimensional space. The rank represents the position of point *j* in the sorted list of nearest neighbors of point *i*. For instance, if *j* is the closest point to *i, r*(*i, j*) would be 1. If it is the second closest, it would be 2, and so forth.

##### Continuity

Continuity [44] is, in a sense, the opposite of trustworthiness. It measures how well the low-dimensional space preserves points that are far apart in the high-dimensional space. This metric helps identify if the algorithm has caused local distortions, where points close in the high-dimensional space become far apart in the low-dimensional space. Continuity is calculted from the following formula:

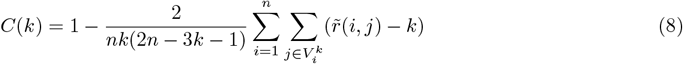

Here 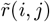 represents the rank of the high-dimensional datapoint *j* according to the pairwise distances between the the high-dimensional datapoints. The variable 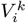 indicates the set of points that are among the *k* nearest neighbors in the high-dimensional space but not in the low-dimensional space

##### Co-Ranking Matrix

The co-ranking matrix, introduced by [45], captures changes in rank distances between high-dimensional and low-dimensional spaces. It provides insights into how well a DR technique preserves local neighborhood relationships.

Let *d*_*ij*_ = *d*(*x*_*i*_, *x*_*j*_) denote the distance between points *x*_*i*_ and *x*_*j*_ in the high-dimensional space, and let 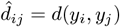 denote the distance between their corresponding projections in the low-dimensional space. We then define the rank of *y*_*j*_ with respect to *y*_*i*_ in the low-dimensional space as:

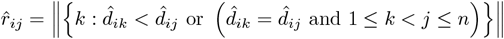

Similarly, the rank in the original high-dimensional space is:

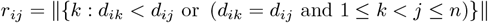

The co-ranking matrix *Q* is then defined by its elements:

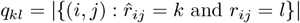

In other words, *q*_*kl*_ counts the number of point pairs whose distance rank was *l* in the high-dimensional space and became *k* in the low-dimensional space.

From the co-ranking matrix, the neighborhood preservation measure *Q*_*NX*_ (*k*) is defined as:

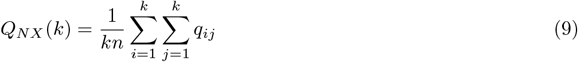

which measures the proportion of points that are among the *k* nearest neighbors in both the original and reduced spaces, normalized so that the maximum value is 1.

##### Implementation Notes

We implemented our own custom function to compute the average Jaccard distance. We used the `ContTrustMeasure` function from version 1.2.2 of the `ProjectionBasedClustering` R package [46] to compute continuity and trustworthiness. Finally, we used the `coranking` and `Q_NX` functions from version 0.2.4 the `coRanking` R package [47] to compute the co-ranking matrices.

### 2.4 Datasets and Pre-processing

In this section, we describe the publicly available biological datasets used in our study. We obtained a total of 4 data sets from public domains for benchmarking DR methods: 2 scRNA, 1 CyTOF, 1 CITE-seq. A summary of the selected datasets is provided in Table 2.

**Table 2:** List of datasets used for benchmarking DR methods.

| Dataset | Type | Tissue | Source | Size |
| --- | --- | --- | --- | --- |
| Patel | scRNA | glioblastoma | GEO | 5498 features, 430 samples |
| Baron | scRNA | pancreas | scRNAseq | 20,125 features, 8569 samples |
| Levine32 | CytoF | bone marrow mononuclear cells | HDCytoData | 265,627 cells, 32 markers |
| PBMC | CITE-seq | peripheral blood mononuclear cells | Zenodo | 161,764 cells, 228 markers |

The primary human glioblastoma scRNA-seq dataset (GSE57872), originally described by Patel et al. [48], was obtained from the NCBI GEO repository [49]. This dataset, recognized as a ‘gold standard’ due to its definitive cell labeling [50], contains 5,948 features and 430 samples. Quality control and visualization were conducted using the SingleCellExperiment and scater packages, following guidelines from the Hemberg Lab’s website [51].

The human pancreas scRNA-seq dataset from Baron et al. [52] is available through the `scRNAseq` R package. This dataset comprises single-cell RNA sequencing data from pancreatic islets of four human donors.

Preprocessing involved standard normalization and scaling procedures using the `Seurat` package, including the identification of highly variable genes (HVGs). Specifically, we applied the “LogNormalize” method, which normalizes feature expression for each cell by dividing raw counts by the total expression, multiplying by a scale factor (default value of 10,000), and applying a log transformation. We then selected 5,000 HVGs, consistent with recommendations from [53] and [43], to capture gene expression variability across cells.

The CyTOF dataset used in this study is a 32-dimensional dataset, which contains protein expression levels for 265,627 cells across 32 protein markers. The data includes cluster labels for 39% of the cells (104,184 out of 265,627), representing 14 manually gated cell populations (clusters), from two individuals. This dataset is sourced from [54]. We used the `HDCytoData` and `SummarizedExperiment` R packages to load it. To normalize the data, we applied an inverse hyperbolic sine (asinh) transformation to the expression values with a cofactor of 5. This step is commonly used for mass cytometry data to compress the dynamic range while preserving the underlying distribution of the data.

We also used the Multimodal PBMC Reference Dataset, which we loaded from Zenodo [55]. This dataset includes data from 161,764 human peripheral blood mononuclear cells (PBMCs) and features antibody panels extending to 228 markers. The dataset is available as a Seurat object, created using Seurat v5. The Seurat object already contains 3000 HVGs, which are used for downstream analysis. We extract the expression data for these genes and save it for later use. Similarly, we extract and save the expression data for the antibody-derived tags (ADT). Since our CITE-seq data includes two distinct assays—gene expression and surface protein measurements—it is essential to integrate these modalities before applying DR. The challenge is to combine the high-dimensional data from both modalities, each offering valuable biological insights, into a single representation that accurately captures the relationships between them. One approach to handling multimodal data is to apply DR separately to each dataset and then align them using a Procrustes transformation. This transformation involves translating, scaling, and rotating the data to align one configuration with another as closely as possible. We use the `procrustes()` function from the `vegan` R package. Procrustes rotation adjusts a matrix to achieve maximum similarity with a target matrix by minimizing the sum of squared differences. The first argument is the target matrix, and the second is the matrix to be rotated. The results will depend on the order of the matrices. We chose the best aligment for each method. We loaded the original subsamples for both RNA and ADT assays, using the same set of cells. While there are other possible approaches to integrating multimodal data, they are beyond the scope of this paper. Notably, `Seurat`, a widely used tool in bioinformatics, primarily offers the weighted nearest neighbor (WNN) approach for multimodal data integration, followed by t-SNE and UMAP for visualization.

The relatively small size of the Patel scRNA dataset allowed us to perform the analysis without the need for downsampling. For the Baron scRNA and Levine32 CyTOF datasets, we randomly subsampled 5% of cells from each cluster and repeated this procedure 10 times. For the PBMC CITE-seq dataset, we randomly selected 1,000 cells from each cluster. We then applied DR algorithms and saved the results (projections and distance matrices) for each repetition and method across all datasets.

## 3 Results

We assessed the performance of 16 DR methods using 4 publicly available datasets: Patel scRNA, Baron scRNA, PBMC CITE-seq and Levine32 CyTOF. A summary of the DR methods with the best alignments for the PBMC CITE-seq dataset, along with the sum of squared differences, can be found in the Supplementary Material. Projections are displayed in Fig. 3, Fig. 4, Fig. 5, and Fig. 6.

**Figure 3.**
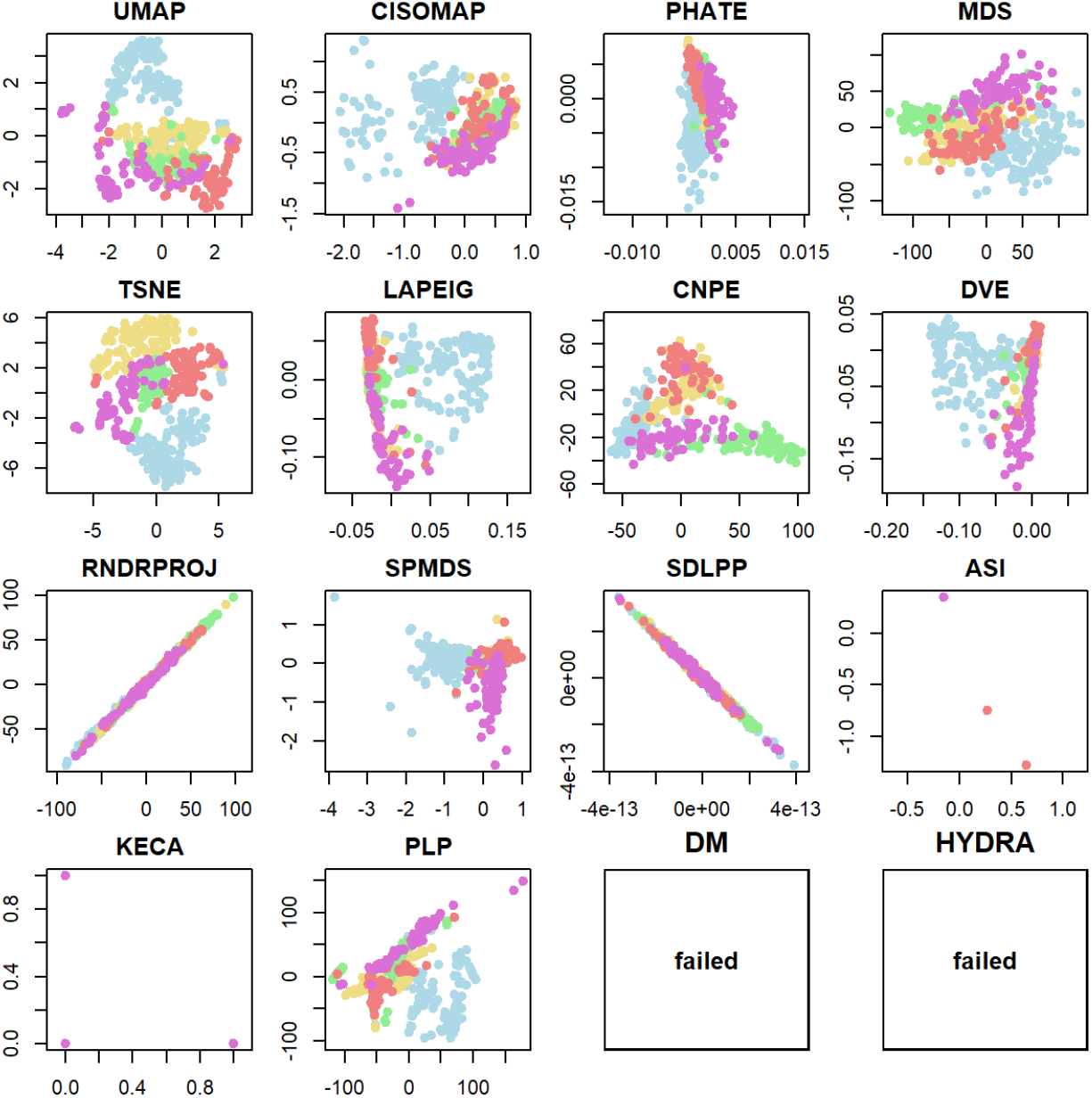
Projections of the Patel scRNA data. DM and HYDRA are excluded due to the generation of NA values. The colors represent clusters corresponding to five patients.

**Figure 4.**
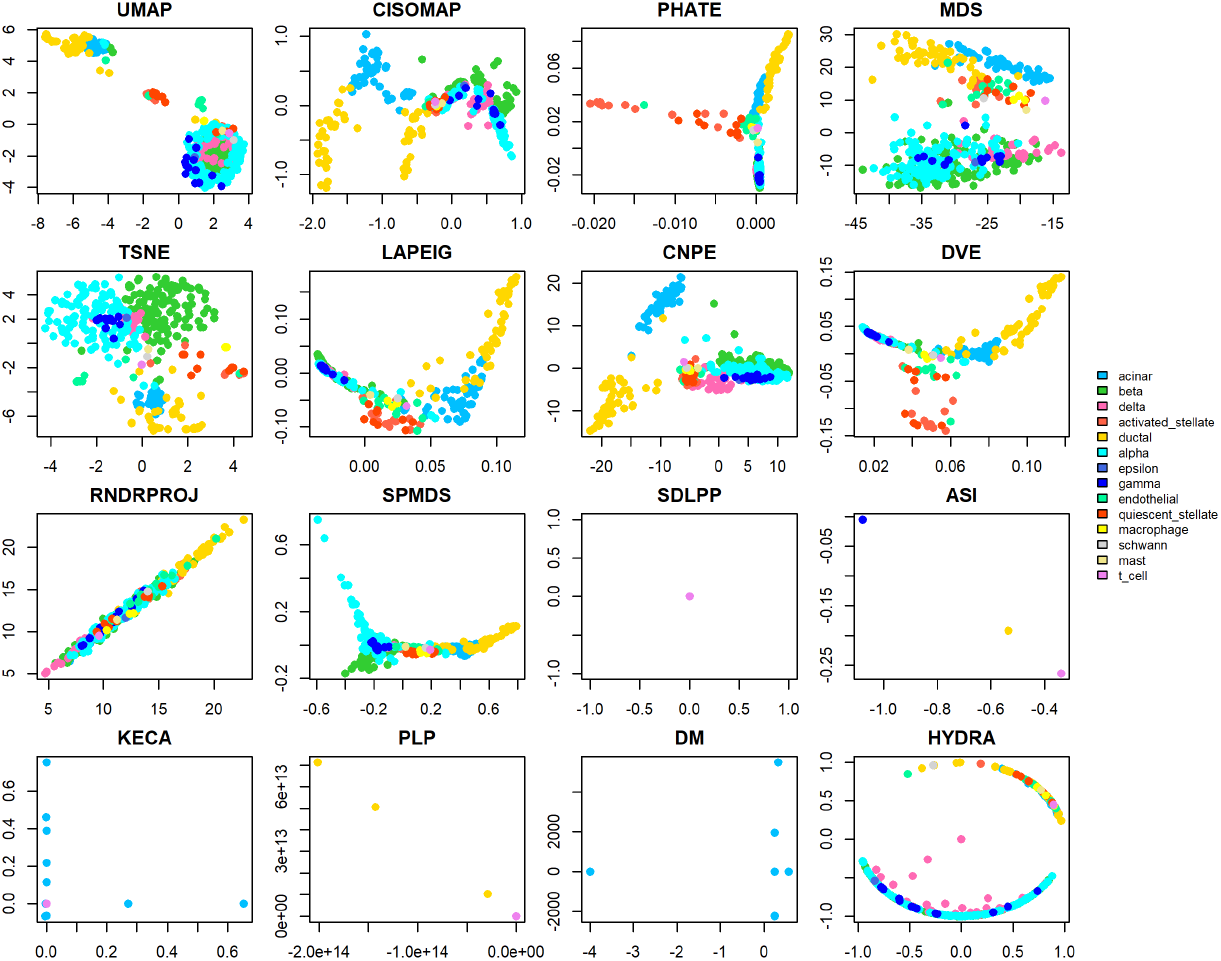
Projections of the Baron scRNA data, subsample 1.

**Figure 5.**
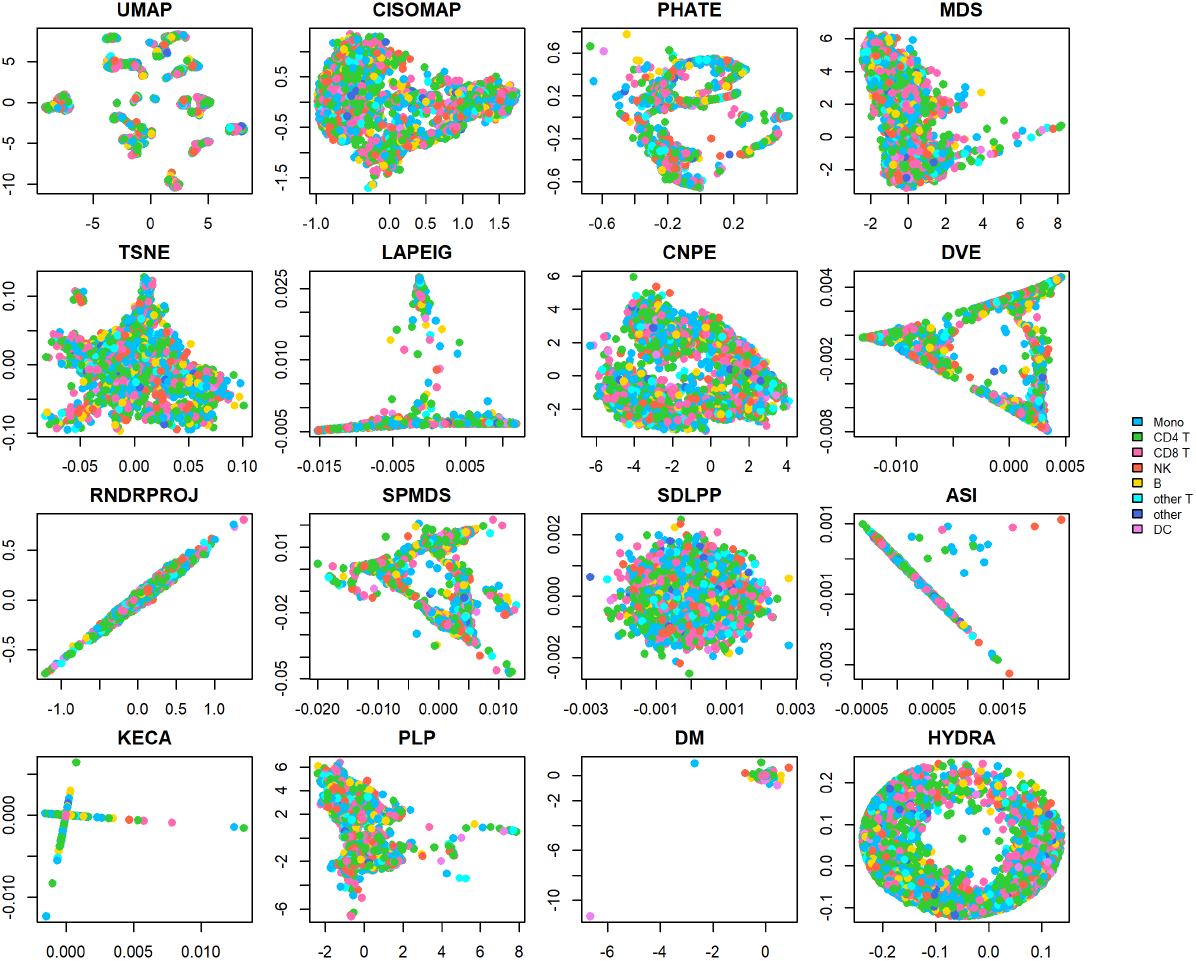
Projections of the procrustes aligmnets of the PBMC CITE-seq dataset.

**Figure 6.**
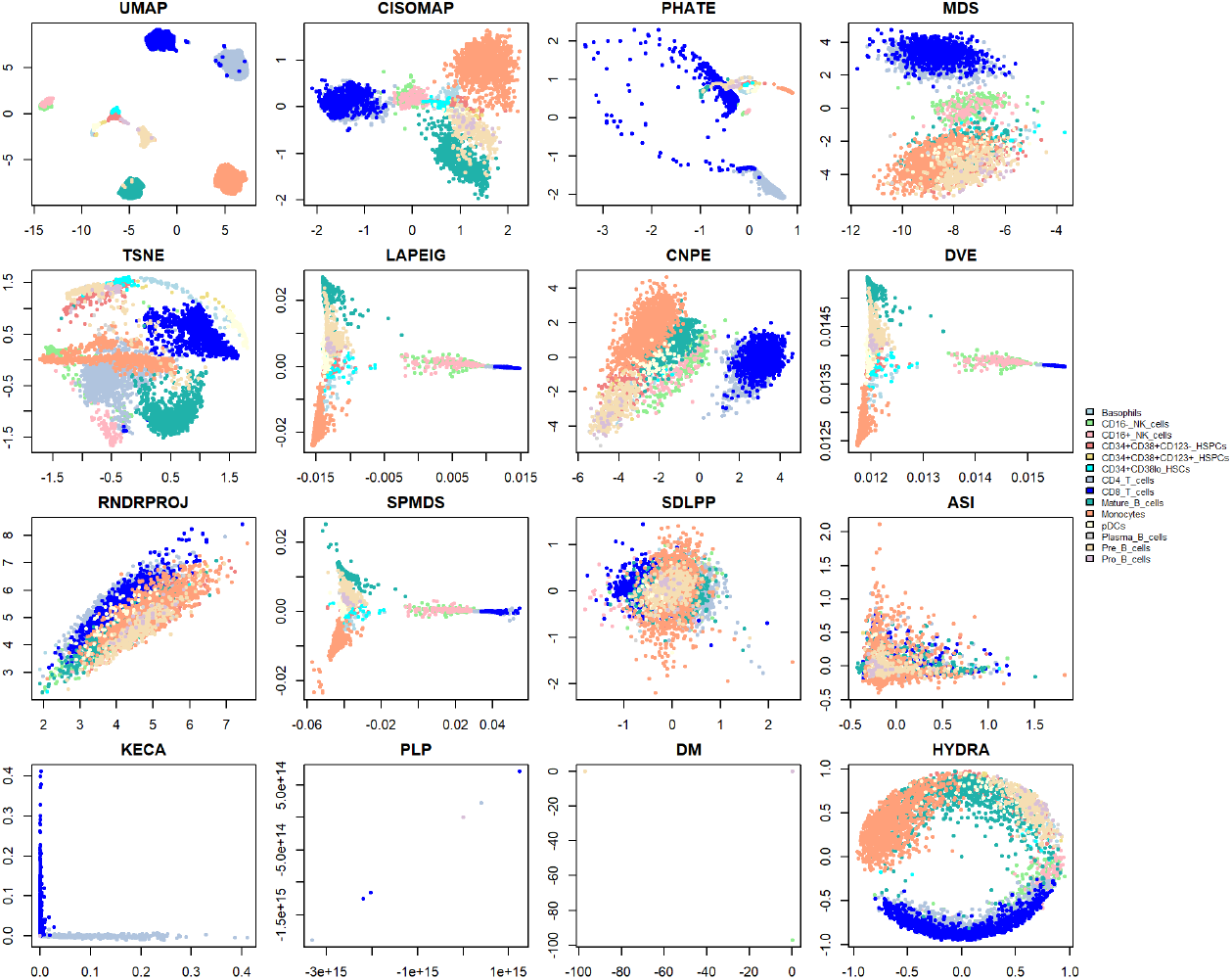
Projections of the Levine32 CyTOF dataset, subsample 1.

### 3.1 Results Based on Pairwise Distance Metrics

The performance of DR methods varied across metrics based on pairwise distance matrices. We present the results in tables (Tables 3 - 8) organized by each metric, with values for different methods displayed as rows and datasets as columns.

**Table 3:**
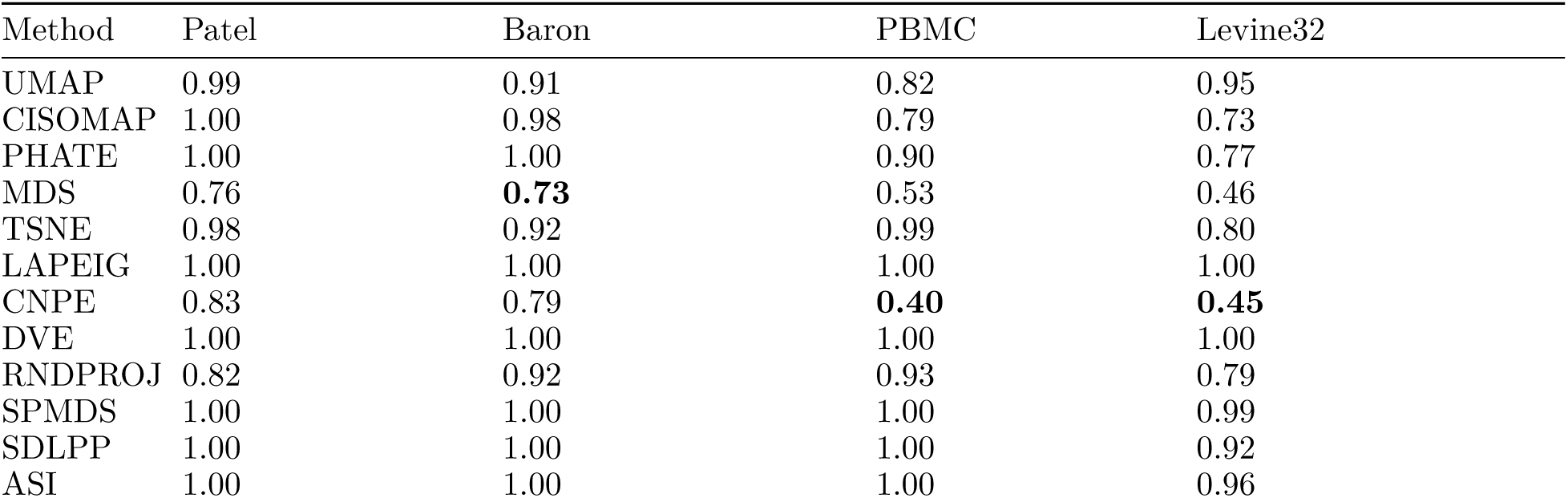
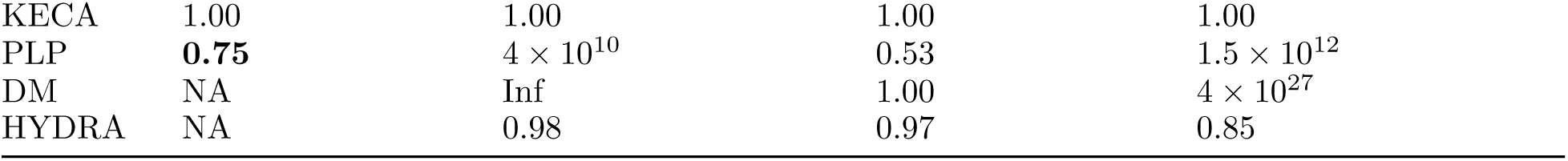
Stress evaluation of dimensionality reduction methods in four datasets (Patel scRNA, Baron scRNA, PBMC CITE-seq, Levine32 CyTOF). NA indicates that stress values could not be calculated for the given method and dataset. Inf indicates that the stress value is infinite. The best value in each column is highlighted in bold.

**Table 4:** Milnor Distortion evaluaton of dimensionality reduction methods in four datasets (Patel scRNA, Baron scRNA, PBMC CITE-seq, Levine32 CyTOF). NA indicates that values could not be calculated for the given method and dataset. Inf indicates that the value is infinite. The best value in each column is highlighted in bold.

| Method | Patel | Baron | PBMC | Levine32 |
| --- | --- | --- | --- | --- |
| UMAP | 7.97 | 8.32 | 11.61 | 10.89 |
| CISOMAP | 7.93 | 8.01 | 10.58 | 9.52 |
| PHATE | 7.69 | 9.82 | 14.76 | 13.83 |
| MDS | **6.83** | **7.02** | 9.69 | 9.38 |
| TSNE | 7.95 | 8.03 | 10.89 | 10.08 |
| LAPEIG | 6.99 | 8.72 | 37.37 | 13.56 |
| CNPE | 7.42 | 8.11 | **9.13** | **9.26** |
| DVE | 9.50 | 9.95 | 11.67 | 14.85 |
| RNDPROJ | 7.93 | 8.44 | 10.26 | 10.17 |
| SPMDS | 7.91 | 7.98 | 12.79 | 12.27 |
| SDLPP | 8.11 | NA | 10.96 | 10.53 |
| ASI | 37.75 | 36.99 | 14.82 | 11.64 |
| KECA | Inf | NA | Inf | 57.82 |
| PLP | Inf | 42.35 | 39.27 | Inf |
| DM | NA | NA | Inf | 42.09 |
| HYDRA | NA | 11.18 | 12.46 | 10.37 |

**Table 5:** M1 distortion evaluation of dimensionality reduction methods in four datasets (Patel scRNA, Baron scRNA, PBMC CITE-seq, Levine32 CyTOF). NA indicates that values could not be calculated for the given method and dataset. Inf indicates that the value is infinite. The best value in each column is highlighted in bold.

| Method | Patel | Baron | PBMC | Levine32 |
| --- | --- | --- | --- | --- |
| UMAP | 1.00 | 0.99 | 1.60 | 0.22 |
| CISOMAP | 1.00 | 1.00 | 0.94 | 0.98 |
| PHATE | 1.00 | 1.00 | 0.99 | 0.97 |
| MDS | 0.92 | **0.62** | 0.54 | **0.15** |
| TSNE | 1.00 | 0.99 | 1.00 | 0.99 |
| LAPEIG | 1.00 | 1.00 | 1.00 | 1.00 |
| CNPE | 0.96 | 0.95 | **0.42** | 0.89 |
| DVE | 1.00 | 1.00 | 1.00 | 1.00 |
| RNDPROJ | 0.95 | 0.89 | 0.99 | 0.55 |
| SPMDS | 1.00 | 1.00 | 1.00 | 1.00 |
| SDLPP | 1.00 | 1.00 | 1.00 | 1.00 |
| ASI | 1.00 | 1.00 | 1.00 | 1.00 |
| KECA | 1.00 | 1.00 | 1.00 | 1.00 |
| PLP | **0.91** | $10^{22}$ | 0.54 | $6 \times 10^{24}$ |
| DM | NA | NA | 1.00 | $5 \times 10^{55}$ |
| HYDRA | NA | 1.00 | 1.00 | 0.99 |

**Table 6:** Sigma distortion evaluation of dimensionality reduction methods in four datasets (Patel scRNA, Baron scRNA, PBMC CITE-seq, Levine32 CyTOF). NA indicates that values could not be calculated for the given method and dataset. Inf indicates that the value is infinite. The best value in each column is highlighted in bold.

| Method | Patel | Baron | PBMC | Levine32 |
| --- | --- | --- | --- | --- |
| UMAP | 0.98 | 0.75 | 3.26 | 3.56 |
| CISOMAP | 0.99 | 0.96 | 0.60 | 0.55 |
| PHATE | 1.00 | 1.00 | 0.77 | 2.09 |
| MDS | **0.56** | **0.52** | 0.66 | 0.37 |
| TSNE | 0.96 | 0.82 | 0.98 | 0.61 |
| LAPEIG | 1.00 | 1.00 | 0.99 | 0.99 |
| CNPE | 0.65 | 0.59 | **0.4** | **0.34** |
| DVE | 1.00 | 1.00 | 1.00 | 1.00 |
| RNDPROJ | 0.64 | 0.79 | 0.82 | 0.62 |
| SPMDS | 0.99 | 0.99 | 0.99 | 0.98 |
| SDLPP | 1.00 | 1.24 | 1.00 | 0.79 |
| ASI | 0.99 | 0.99 | 1.00 | 0.86 |
| KECA | 0.99 | 1.00 | 1.00 | 0.97 |
| PLP | 0.56 | $7 \times 10^{25}$ | 0.69 | $2.6 \times 10^{31}$ |
| DM | NA | NA | 1331.97 | $10^{63}$ |
| HYDRA | NA | 0.96 | 0.92 | 0.70 |

**Table 7:** Spearman Rho evaluation of dimensionality reduction methods in four datasets (Patel scRNA, Baron scRNA, PBMC CITE-seq, Levine32 CyTOF). NA indicates that values could not be calculated for the given method and dataset. The best value in each column is highlighted in bold.

| Method | Patel | Baron | PBMC | Levine32 |
| --- | --- | --- | --- | --- |
| UMAP | 0.61 | 0.43 | 0.36 | 0.63 |
| CISOMAP | 0.6 | 0.35 | 0.7 | **0.84** |
| PHATE | 0.61 | 0.52 | 0.35 | 0.26 |
| MDS | 0.49 | 0.45 | 0.76 | 0.81 |
| TSNE | 0.62 | **0.57** | 0.01 | 0.38 |
| LAPEIG | 0.7 | 0.46 | 0.45 | 0.67 |
| CNPE | 0.14 | 0.37 | **0.83** | 0.83 |
| DVE | **0.76** | **0.57** | 0.58 | 0.68 |
| RNDPROJ | 0.35 | 0.12 | 0.39 | 0.34 |
| SPMDS | 0.62 | 0.49 | 0.79 | 0.79 |
| SDLPP | 0.04 | 0.02 | -0.02 | 0.1 |
| ASI | 0.45 | 0.41 | 0.41 | 0.07 |
| KECA | -0.03 | -0.19 | -0.09 | -0.02 |
| PLP | 0.43 | 0.45 | 0.74 | 0.76 |
| DM | NA | NA | -0.16 | 0.75 |
| HYDRA | NA | 0.49 | 0.59 | 0.79 |

**Table 8:** EMD evaluation of dimensionality reduction methods in four datasets (Patel scRNA, Baron scRNA, PBMC CITE-seq, Levine32 CyTOF). NA indicates that values could not be calculated for the given method and dataset. The best value in each column is highlighted in bold.

| Method | Patel | Baron | PBMC | Levine32 |
| --- | --- | --- | --- | --- |
| UMAP | 581 | 1.94 | 8 | 8.22 |
| CISOMAP | 582 | 1.82 | 9 | 12.13 |
| PHATE | 584 | 1.77 | 11 | 11.53 |
| MDS | 468 | **1.74** | 2 | 6.85 |
| TSNE | 605 | 1.95 | 10 | 11.43 |
| LAPEIG | 584 | 1.77 | 10 | 12.64 |
| CNPE | 523 | 1.83 | 2 | 6.43 |
| DVE | 584 | 1.77 | 14 | 10.86 |
| RNDPROJ | 510 | 1.91 | 11 | 11.06 |
| SPMDS | 582 | 1.8 | 16 | 13.97 |
| SDLPP | 584 | **1.74** | 8 | 12.36 |
| ASI | 584 | 1.87 | 8 | 12.16 |
| KECA | 582 | 1.76 | 13 | 15.43 |
| PLP | **459** | 1.8 | **1** | **6.18** |
| DM | NA | NA | 7 | $6 \times 10^{12}$ |
| HYDRA | NA | 1.82 | 12 | 12.17 |

A consistent observation is the superior performance of both CNPE and MDS. MDS excels across various metrics, particularly in scRNA datasets. Similarly, CNPE demonstrates robust performance, especially in the PBMC CITE-seq and Levine32 CyTOF datasets. The performance of the PLP method is highly variable. While PLP stands out as the best method for the EMD metric, outperforming others in 3 out of 4 datasets (Patel scRNA, PBMC CITE-seq, and Levine32 CyTOF), it frequently produces extreme off-scale or undefined values. Methods commonly used for visualization of biological dataset, such as TSNE and UMAP, generally show poor performance. Although TSNE preserves correlations well (e.g., in the Baron scRNA dataset), both methods perform poorly in other metrics based on pairwise distance matrices. Notably, UMAP produces M1 and sigma distortion values exceeding 1 in the PBMC dataset. DM, KECA, and HYDRA consistently underperform across all metrics.

### 3.2 Results Based on Neighborhood Preservation Metrics

Next, we outline the results for metrics focused on neighborhood preservation. Table 9 summarizes the Average Jaccard Distance (AJD) for various DR methods.

**Table 9:** Average Jaccard distance in four datasets (Patel scRNA, Baron scRNA, PBMC CITE-seq, Levine32 CyTOF). For the Patel scRNA, Baron scRNA, and Levine32 CyTOF datasets,k = 100; for the PBMC CITE-seq dataset, k = 300. NA indicates that values could not be calculated for the given method and dataset. The best value in each column is highlighted in bold, except for the Baron scRNA dataset, where all methods show poor performance.

| Method | Patel | Baron | PBMC | Levine32 |
| --- | --- | --- | --- | --- |
| UMAP | 0.64 | 0.99 | **0.73** | **0.48** |
| CISOMAP | 0.69 | 0.99 | 0.75 | 0.75 |
| PHATE | 0.73 | 0.99 | 0.79 | 0.52 |
| MDS | 0.67 | 0.99 | 0.82 | 0.96 |
| TSNE | **0.61** | 0.99 | 0.92 | 0.59 |
| LAPEIG | 0.64 | 0.99 | 0.87 | 0.87 |
| CNPE | 0.7 | 0.99 | 0.77 | 0.95 |
| DVE | 0.62 | 0.99 | 0.75 | 0.88 |
| RNDRPROJ | 0.76 | 0.99 | 0.93 | 0.98 |
| SPMDS | 0.69 | 0.99 | 0.75 | 0.86 |
| SDLPP | 0.86 | 0.99 | 0.98 | 0.99 |
| ASI | 0.72 | 0.99 | 0.96 | 1 |
| KECA | 0.92 | 0.99 | 0.96 | 0.97 |
| PLP | 0.73 | 0.99 | 0.82 | 0.94 |
| DM | NA | NA | 0.95 | 0.53 |
| HYDRA | NA | 0.99 | 0.93 | 0.97 |

**Table 10:** Summary of dimensionality reduction methods with best alignments and the sum of squared differences (ssSum) Values.

| Method | Best Alignment | ssSum |
| --- | --- | --- |
| UMAP | RNA to ADT | 244439.500 |
| CISOMAP | ADT to RNA | 4037.916 |
| PHATE | RNA to ADT | 1842.225 |
| MDS | ADT to RNA | 40186.160 |
| tSNE | RNA to ADT | 44.720 |
| LAPEIG | RNA to ADT | 0.407 |
| CNPE | ADT to RNA | 52291.840 |
| DVE | ADT to RNA | 0.432 |
| RNDPROJ | ADT to RNA | 6947.362 |
| SPMDS | ADT to RNA | 5.814 |
| SDLPP | RNA to ADT | 48.297 |
| ASI | RNA to ADT | 3.906 |
| KECA | RNA to ADT | 1.987 |
| PLP | ADT to RNA | 40994.890 |
| DM | ADT to RNA | 23905879.000 |
| HYDRA | RNA to ADT | 2875.112 |

Across all analyzed datasets, TSNE and UMAP stand out as strong performers in preserving local neighborhood structures. However, other methods also exhibit potential, depending on the specific dataset and context. For example, LAPEIG delivers promising results in the Patel scRNA dataset, while DVE, CISOMAP, and SPMDS perform well in the PBMC CITE-seq dataset. In the Levine32 CyTOF dataset, PHATE closely follows UMAP in performance.

Surprisingly, the average trustworthiness (Fig 7) and continuity graphs (Fig 8) reveal that all methods exhibit low trustworthiness values (a score above 0.80 is considered to be a “good” result). This could be due to the fact that we did not test other values of *k*, which might have influenced the results. Among the evaluated methods, SDLPP consistently demonstrates superior performance, achieving the highest trustworthiness and continuity scores across most datasets.

**Figure 7.**
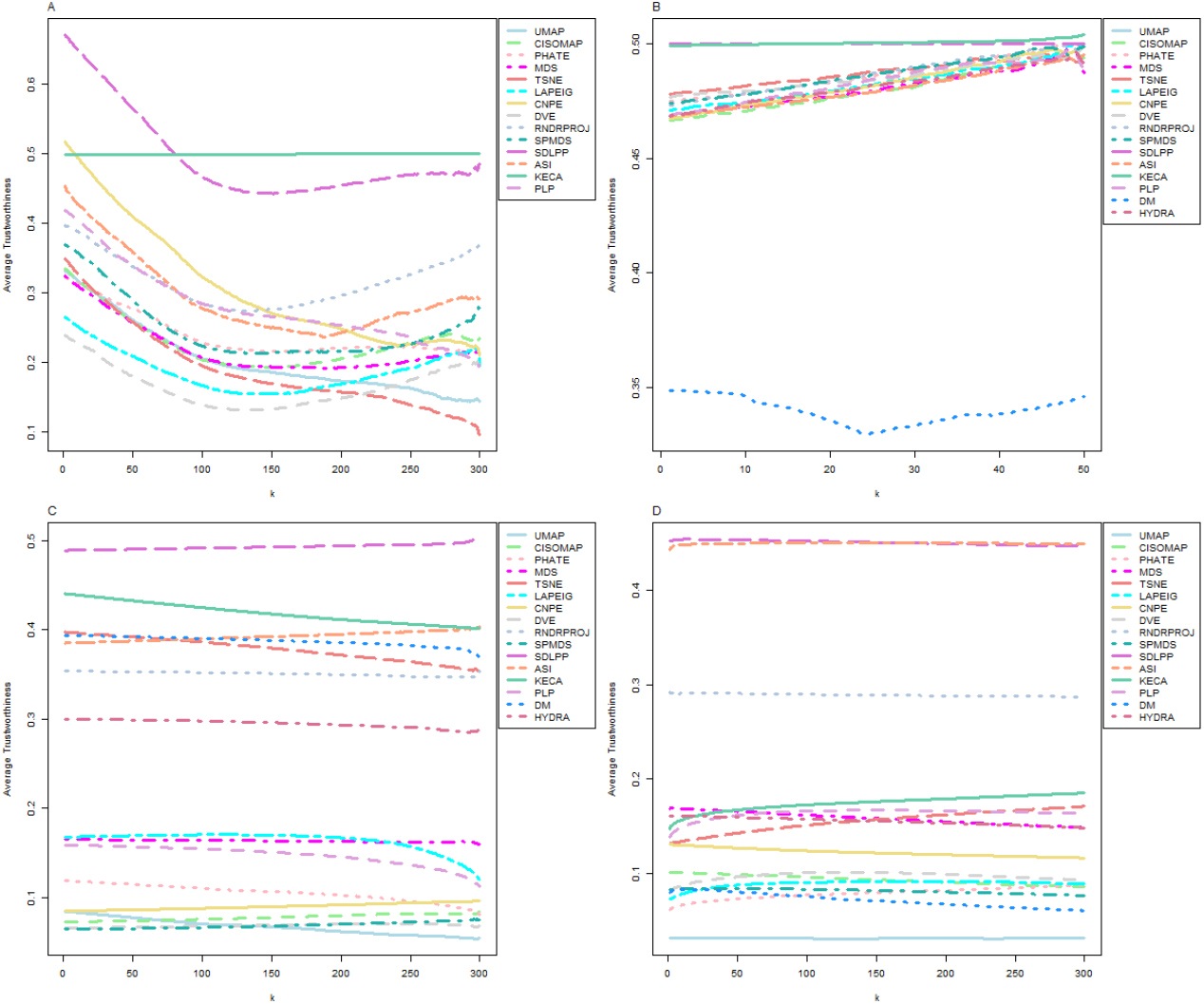
Average trustworthiness for each dimensionality reduction method applied to: (A) Patel scRNA dataset; (B) Baron scRNA dataset; (C) PBMC CITE-seq dataset; (D) Levine32 CyTOF dataset.

**Figure 8.**
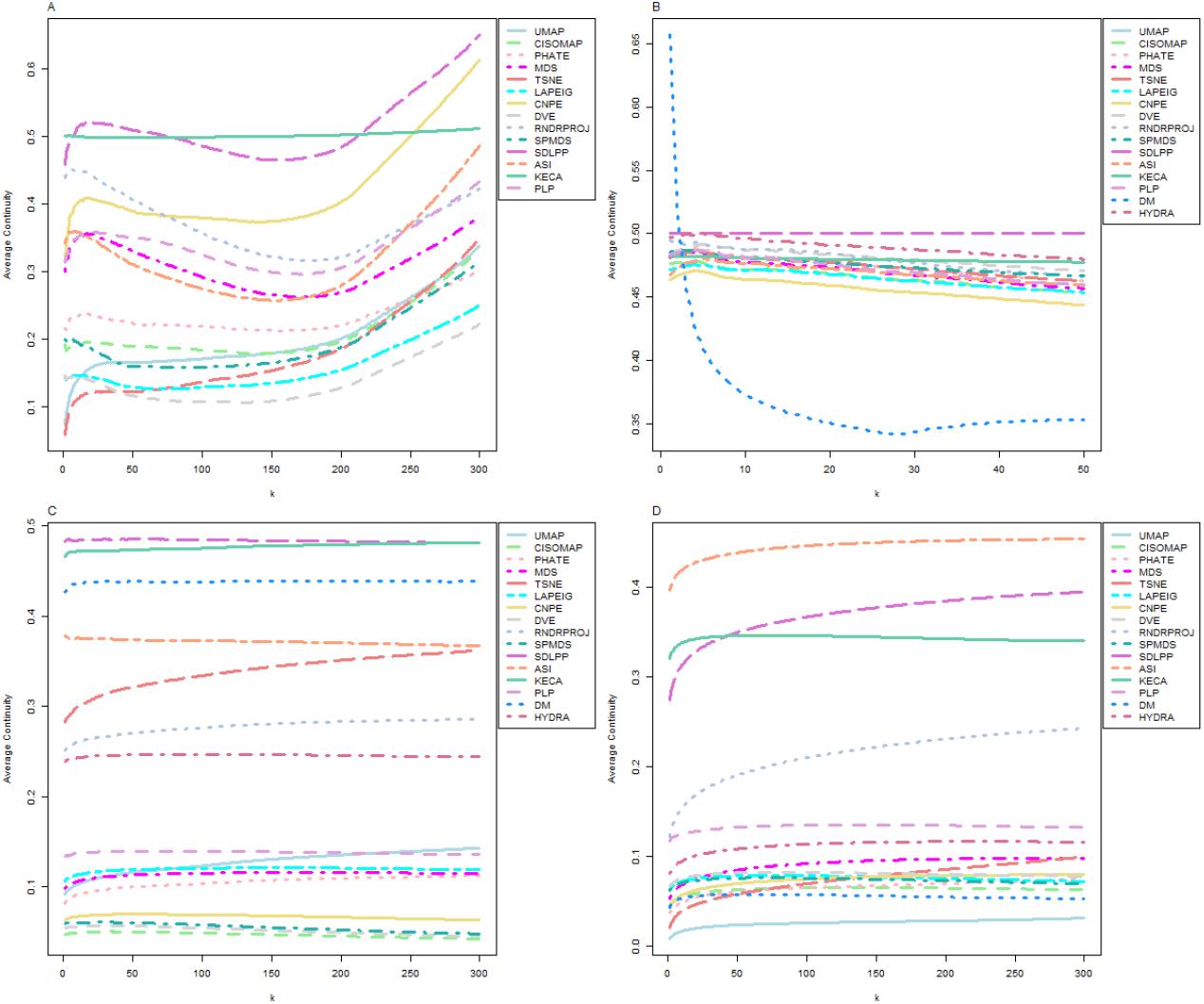
Average continuity for each dimensionality reduction method applied to: (A) Patel scRNA dataset; (B) Baron scRNA dataset; (C) PBMC CITE-seq dataset; (D) Levine32 CyTOF dataset.

Then we turn our attention to the quality curves derived from the co-ranking matrices (Fig 9). Clear performance patterns emerge: MDS and CNPE stand out as the most reliable methods, while TSNE (Fig 9 (A) and (B)) and UMAP (C) excels only at capturing relatively small neighborhoods. In contrast, methods like SDLPP and KECA consistently perform poorly across all scenarios, highlighting their limitations in preserving neighborhood structure.

**Figure 9.**
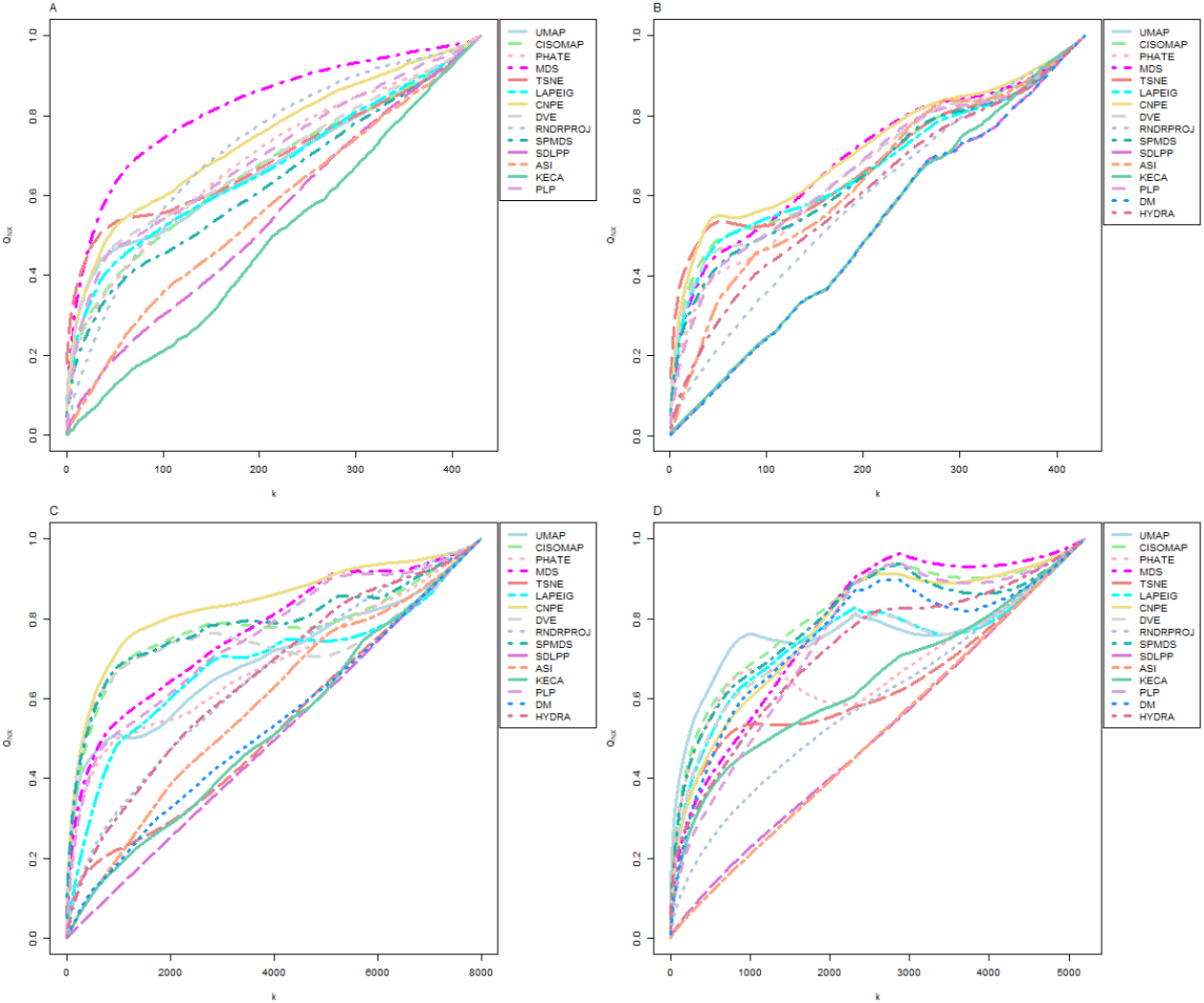
Quality curve for each dimensionality reduction method applied to: (A) Patel scRNA dataset; (B) Baron scRNA dataset; (C) PBMC CITE-seq dataset; (D) Levine32 CyTOF dataset.

## 4 Discussion

In this work, we evaluate 16 DR algorithms on three main types of biological data from two perspectives: metrics based on pairwise distance matrices and metrics based on neighborhood preservation. The data used includes scRNA, CYTOF, and CITE-seq. Our analysis provides insights into the strengths and weaknesses of each method.

The projections revealed several issues. Some algorithms, such as DM, HYDRA, and PLP, encountered NA values, likely indicating numerical instability. ASI exhibited a tendency for points to collapse into identical coordinates, suggesting that the stopping criteria may not have been appropriately configured. KECA generated projections with very few non-zero values. RNDRPROJ resulted in points forming a straight line with a positive slope, while SDLPP produced a straight line with a negative slope in some projections.

Overall, we observed that CNPE and MDS consistently achieved the best results across multiple quality metrics, including stress, Milnor distortion, and M1 distortion, across the majority of the analyzed biological data. Additionally, CNPE and MDS were top performers according to the quality curves derived from co-ranking matrices, which quantify the consistency of local neighborhoods across dimensions. In contrast widely used DR methods such as TSNE and UMAP have difficulty capturing the global structure.

Before concluding this manuscript, we must acknowledge several limitations of the present study. First, when running DR algorithms from the `Rdimtools` R package, we relied solely on the default hyperparameter values. Additionally, the performance of these algorithms may be heavily influenced by the dataset size and the chosen downsampling method. The CITE-seq dataset was preprocessed using the Procrustes alignment strategy, and we did not explore other possible integration techniques. Moreover, we did not investigate the potential of combining multiple DR algorithms to assess whether integrating different methods could address individual weaknesses and improve overall performance. Finally, further analysis with additional datasets would likely enhance the robustness and generalizability of the results.

## 5 Conslusion

In conclusion, CNPE shows great promise as a valuable tool for scholars and bioinformaticians working on dimensionality reduction of biological data. While we verified that MDS is a strong method, CNPE serves as an additional, highly effective tool that enhances the dimensionality reduction process, offering further opportunities for advancing research in this field.

## 6 Supplementary Material

## 7 Acknowledgements

## Notes

### Competing Interest Statement

The authors have declared no competing interest.

